# Robust Microfabrication of Highly Parallelized Three-Dimensional Microfluidics on Silicon

**DOI:** 10.1101/625277

**Authors:** Sagar Yadavali, Daeyeon Lee, David Issadore

## Abstract

We present a new, robust three dimensional microfabrication method for highly parallel microfluidics, to improve the throughput of on-chip material synthesis by allowing parallel and simultaneous operation of many replicate devices on a single chip. Recently, parallelized microfluidic chips fabricated in Silicon and glass have been developed to increase the throughput of microfluidic materials synthesis to an industrially relevant scale. These parallelized microfluidic chips require large arrays (> 10,000) of Through Silicon Vias (TSVs) to deliver fluid from delivery channels to the parallelized devices. Ideally, these TSVs should have a small footprint to allow a high density of features to be packed into a single chip, have channels on both sides of the wafer, and at the same time minimize debris generation and wafer warping to enable permanent bonding of the device to glass. Because of these requirements and challenges, previous approaches cannot be easily applied to produce three dimensional microfluidic chips with a large array of TSVs. To address these issues, in this paper we report a fabrication strategy for the robust fabrication of three-dimensional Silicon microfluidic chips consisting of a dense array of TSVs, designed specifically for highly parallelized microfluidics. In particular, we have developed a two-layer TSV design that allows small diameter vias (*d* < 20 *µ*m) without sacrificing the mechanical stability of the chip and a patterned SiO_2_ etch-stop layer to replace the use of carrier wafers in Deep Reactive Ion Etching (DRIE). Our microfabrication strategy allows >50,000 (*d* = 15 *µ*m) TSVs to be fabricated on a single 4” wafer, using only conventional semiconductor fabrication equipment, with 100% yield (*M* = 16 chips) compared to 30% using previous approaches. We demonstrated the utility of these fabrication strategies by developing a chip that incorporates 20,160 flow focusing droplet generators onto a single 4” Silicon wafer, representing a 100% increase in the total number of droplet generators than previously reported. To demonstrate the utility of this chip for generating pharmaceutical microparticle formulations, we generated 5–9 µm polycaprolactone particles with a CV <5% at a rate as high as 60 g/hr (> 1 trillion particles / hour).

## Introduction

In many sub-fields of microfluidics, parallelization – the placing of many replicate devices that operate in parallel onto a single chip-has been a successful strategy to increase the throughput of otherwise slow processes.^1-16^ Parallelization has been used with particular success to increase the production rate of microfluidic generated materials to the scale required for economic commercial use, including nanomaterials, microparticles, and a variety of single and multiple emulsions.^1-12^ In particular, microfluidic generated micro and nanoparticles have shown excellent pharmacokinetic properties, superior control over drug release rates, long term stable formulations and higher drug encapsulation efficiencies compared to conventional approaches, such as ball milling.^13^ This approach has also been applied successfully to increase the throughput of micro-sensors to detect cells and molecular markers^14-17^ and to perform digital droplet based assays.^18,19^ To deliver fluid to and collect fluid from many parallel microdevices on a single chip, three-dimensional microfabrication strategies have been used to incorporate a layer of delivery channels and vias that connect these delivery channels to microfluidic devices in a layer below.^1,5,9,10^ Three-dimensional fabrication strategies have been developed for both polymer (PDMS, perfluoropoly-ether-polyethylene glycol, etc.)^5,11^ and for Silicon^9,20,21^ based devices. Using these fabrication methods, architectures have been developed that make it possible to operate many microfluidic droplet generators in parallel.^22-23^ Although Silicon devices are significantly more expensive than polymer devices, they have several key advantages that motivate their use, in particular for the generation of micro- and nanomaterials. Importantly, Silicon devices can operate at high pressure *P*_max_ > 1000 PSI and high temperature *T*_max_> 500°C, use solvents useful for material synthesis but that are incompatible with polymer devices, and can be fabricated with significantly less variance in device dimensions than soft-lithography based devices, resulting in more monodispersed materials.^6,20^

The microfabrication of Silicon to create highly parallelized microfluidic devices has several fabrication challenges, which have not been adequately addressed in prior studies^20^ and which must be addressed to produce high performing chips with high yield. Conventional TSV approaches cannot be easily applied to these architectures **1.** because of the small foot-print of these TSVs (d < 20 *µ*m), necessary to allow many microfluidic devices to be packed onto a single wafer, **2.** because of the requirement of these chips for microfabricated features on both sides of the wafer, and **3.** because of the stringent requirement for minimal debris and wafer warping such that the device can be permanently bonded to glass. We describe a set of microfabrication strategies specifically for parallelized microfluidics in Silicon and glass (**Fig. 1a**). By incorporating the microfabrication strategies described in this paper, we achieved a device fabrication yield of 100%(*M* = 16 wafers) using conventional semiconductor fabrication modalities compared to 30% using previous approaches.^20^ The two key innovations in this paper to address these issues, and which make use of conventional MEMS fabrication modalities, are: **1**. We have developed a technique to through-etch the Si to produce vias that are compatible with the specific requirements of parallelized microfluidics, *i.e.* the requirement for DRIE features on both sides of the wafer and the requirement for minimal debris and wafer warping. To accomplish this goal, we replace the conventionally used carrier wafer with a stress relieved SiO_2_ membrane as an etch-stop for DRIE. **2.** We have implemented trenches in our delivery channel, which allow the tradeoff between the diameter of the vias and the mechanical stability of the wafer to be eliminated, allowing the fabrication of small footprint devices to allow many replicate devices to be incorporated onto a single chip.

**Figure 1.**
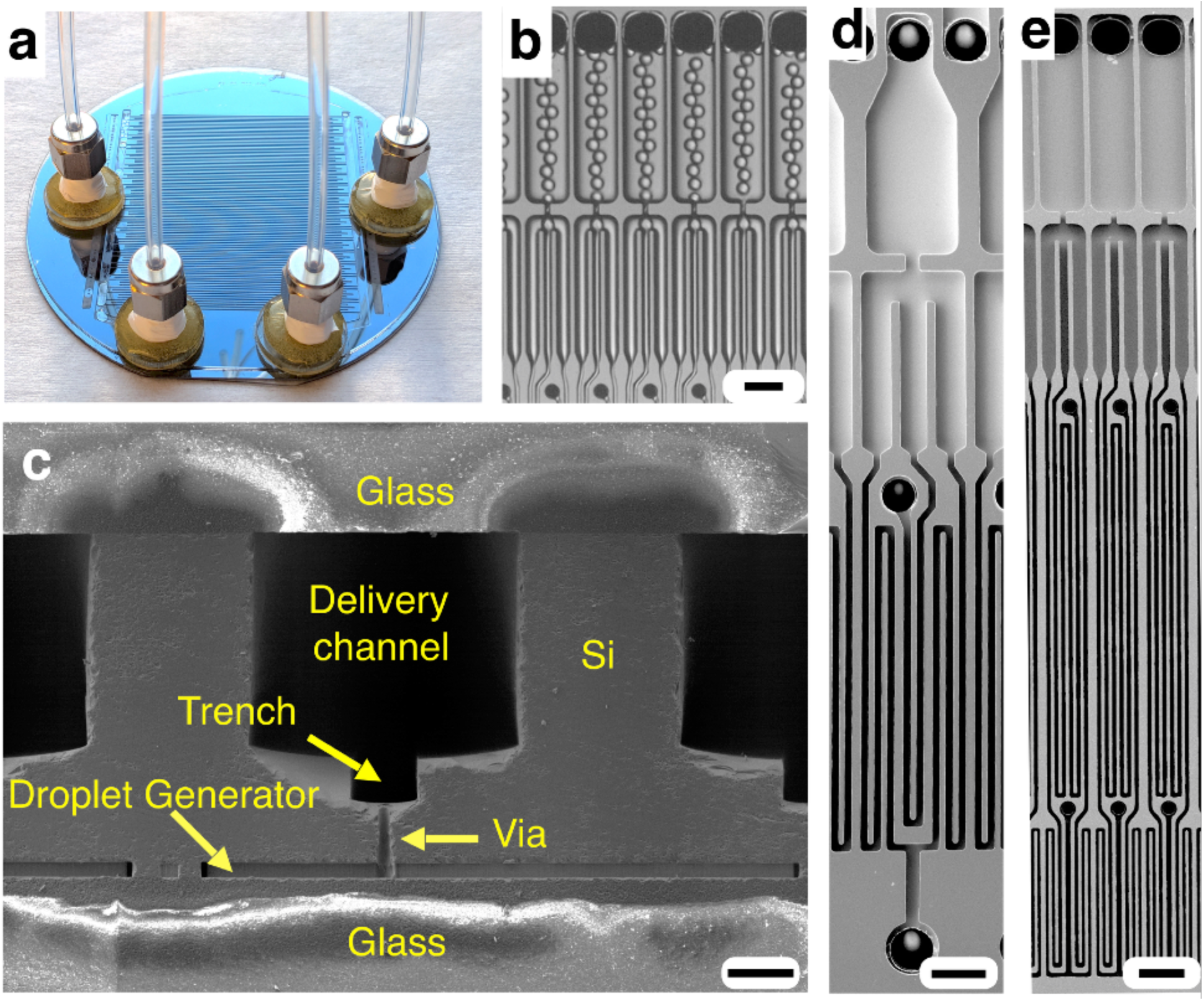
**a.** A photograph of the 20k-VLSDI chip (4” wafer), which consists of 20,160 (565 × 36) flow focusing generators (FFGs). **b.** An optical micrograph of individual droplet generators producing droplets. Scale bar 80 *µ*m. **c.** An SEM micrograph of the cross section of the chip. Scale bar 115 *µ*m. **d.** The FFGs used in our previous 10k-VLSDI chip^19^. Scale bar 80 *µ*m. **e.** FFGs used in this work, where the device footprint has been scaled down by a factor of two compared to the prior work^19^. Scale bar 80 *µ*m.

To demonstrate the utility of these fabrication strategies, we have developed a Very Large Scale Droplet Integration (VLSDI) chip that incorporates 20,160 flow focusing droplet generators onto a single 4” Silicon wafer, representing a 100% increase in the total number of droplet generators on a single chip than has previously been reported (**Fig. 1b**).^1,2,4-6,9,20^ Using these generators that are connected in a ladder geometry with only one set of inlets and outlets, we generated 1 trillion monodispersed droplets / hour with a CV < 5% for diameters ranging from 21-28 *µ*m. We have also generated monodispersed polycaprolactone (PCL) solid microparticles (*d*_p_ = 5.3 – 9.0 *µ*m) with a coefficient of variation CV <5% at a production rate as high as 60 grams/hr. We believe our fabrication strategy can be widely used to enhance the reliability of microfabrication of three-dimensional Silicon and glass microfluidic chips for a variety of applications.

## Results

### The VLSDI Chip

Our 3D fabricated parallelized chips is built using a 4” diameter 500 *µ*m thick double side polished Si wafer (University Wafers, Part 1095). The chip consists of 370 *µ*m height delivery channels on one side of the wafer and includes a trench that has a depth of 75 *µ*m. On the other side of the wafer, there are flow focusing droplet generators with a height *h* = 24 *µ*m. Connecting these two layers of microfluidics are vias with a diameter *d* = 15 *µ*m and a height *h* = 85 *µ*m. Both sides of the wafer are permanently bonded with Borofloat 33 glass (University wafers, Part 517) using anodic bonding to encapsulate the channels (**Fig. 1c**). In this work, we have increased the total number of droplet generators that can be incorporated onto a single wafer by reducing the size of the vias, resulting in a footprint (80 *µ*m × 1.6 mm) for each droplet generator that is 50% smaller than that reported in our previous work (**Fig. 1d,e**).^20^

### Microfabrication Challenges

The microfabrication of VLSDI chips in Silicon is made particularly challenging because of the Through Silicon Vias (TSVs) required to create the connections between the delivery channels and the droplet generators.^24^ Although there is a well developed literature on TSVs for MEMS applications and for CMOS chips,^24-29^ VLSDI chips have several unique requirements that warrant special consideration. Unlike many through-etching applications, parallelized microfluidic chips require microfabricated patterns on both sides of the chip, making many existing through-etching techniques not easy to implement, i.e. the use of mechanical polishing to expose TSVs.^30^ Moreover, parallelized microfluidic chips must be anodically bonded with glass to encapsulate the microfluidic devices, making the fabrication process sensitive to wafer warping and to contamination from debris. Finally, from a design perspective, in VLSDI chips it is advantageous to pack as many parallelized chips onto a wafer as possible. Therefore it is important that the vias have the smallest possible footprint. This requirement makes techniques such as anisotropic wet etching (e.g. potassium hydroxide KOH, tetramethylammonium hydroxide TMAH) unfavorable, due to their angled sidewalls.^31^ Finally, VLSDI chips often require high aspect ratio features (height / width ∼ 5), motivating the use of Deep Reactive Ion Etch (DRIE) for the etching of the microfluidic channels.

### Etching of Through Silicon Vias (TSVs)

Through-etching using DRIE is challenging because in a typical DRIE Bosch process the backside of the wafer is kept at a positive pressure to keep the wafer at a low temperature using He gas, and the frontside is maintained under a vacuum necessary for reactive ion etching.(**Fig. SI 7**) When a TSV punches through a Silicon wafer it connects the positive pressure and vacuum, halting the etching process. The etch quality, uniformity across the wafer, and photoresist selectivity depends on the wafer temperature, and as such the He cooling is necessary. This problem is particularly pertinent for parallelized microfluidics, because of the variety of the diameter of TSVs used. Larger diameter TSVs etch faster, and therefore break through the Silicon before the smaller ones (**Fig. SI 8**). Conventionally, chips are often bonded to a carrier wafer using a temporary adhesive like crystal bond to facilitate TSV etching(**Fig. 2a**).^32^ However, it can be challenging to remove all air pockets between the two wafers, which leads to wafer breakage in the DRIE’s vacuum (**Fig. 2b**). In addition, even if the wafers do not break, the presence of small air pockets between the carrier wafer and the VLSDI can lead to nonuniform etching on the wafer. (**Fig. 2c**). The yield of our VLSDI chip when we used a carrier wafer was only 30% (**Fig. 2d**). By combining our two innovations, i.e our trench technique to achieve small vias and our stress relieved oxide backing layer, we have achieved a yield of 100% (*N* = 20,160 devices per chip, *M* = 16 chips).

**Figure 2.**
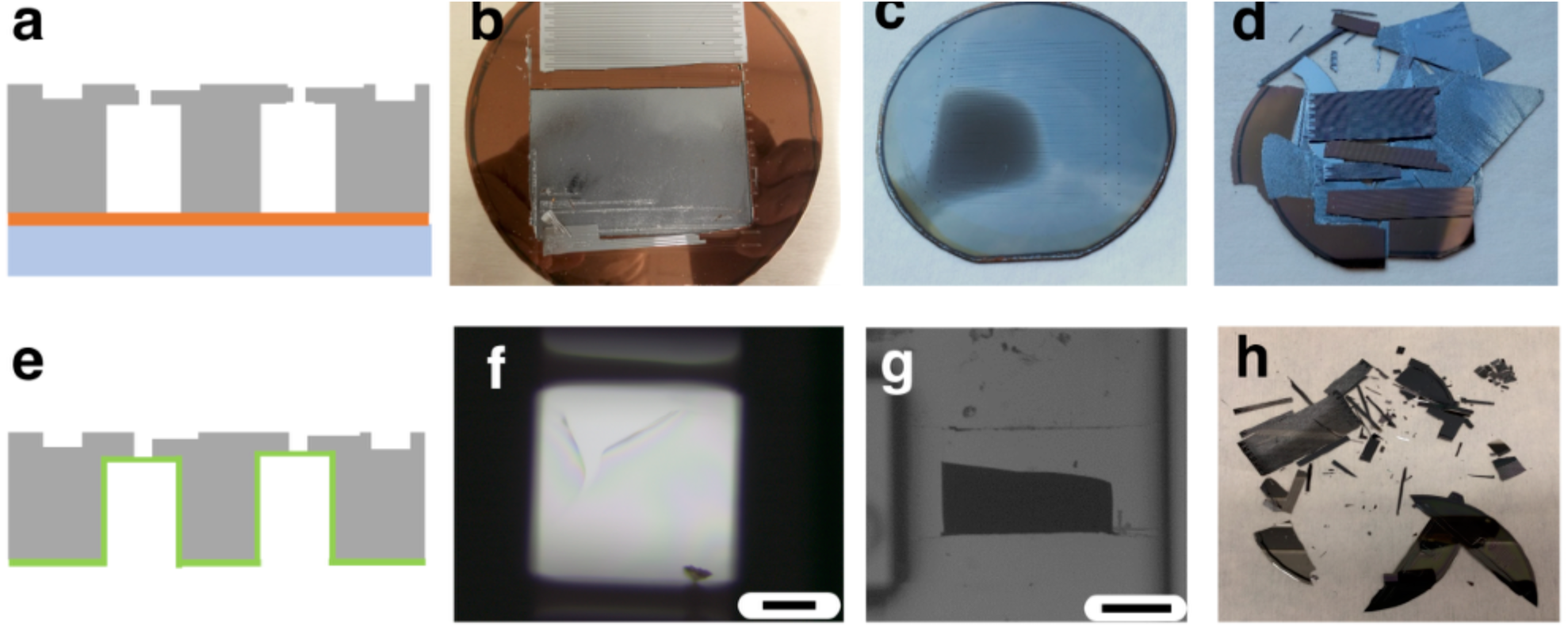
Role of fabrication strategies on the structural integrity and etch quality of 20k-VLSDI chips using conventional approaches. **a.** Schematic showing conventional TSV etching using a carrier wafer. **b.** A photograph shows broken wafer in DRIE chamber that is bonded to carrier wafer using crystal bond adhesive, due to trapped air pockets between the wafers. **c.** A photograph shows non uniform etching of TSVs due to non-uniform temperature across the wafer when using a carrier wafer. **d.** A photograph shows a wafer that broke while being removed from the carrier wafer. The yield using a carrier wafer was 30%. **e.** Schematic showing our technique to replace the carrier wafer with a PECVD SiO_2_ etch-stop, but where we do not include stress relief for the oxide or trenches in the delivery channel. **f.** Micrograph shows broken oxide layer at underpass *via* positions, due to stress relief features not being included. Scale bar 100 *µ*m. **g.** Micrograph shows broken wafer at the intersection of underpass and supply channel positions due to deep etching of delivery channels (*h* = 460 *µ*m), due to trenches not being included. Scale bar 500 *µ*m. **h.** Photograph shows a 3D etched wafer broken during drying with a Nitrogen gun, due to trenches not being included. The yield using this technique was 50%.

In our aproach, instead of using a carrier wafer we instead use a Chemical Vapor Deposition (CVD) grown oxide layer as an etch stop (**Fig. 2e**). A 6 *µ*m thick layer of CVD grown SiO_2_ is deposited on to the delivery channels side of the wafer before the vias are etched. The use of an SiO_2_ back-layer obviates the need for the temporary adhesives and associated problems with uniformity of temperature across the wafer during processing. To avoid mechanical stress on the wafer from the 6 *µ*m thick SiO_2_ layer, which can lead to wrinkling of the SiO_2_ membrane and breaking of the Si wafer (**Fig. 2f**), the SiO_2_ layer is lithographically patterned to relieve the mechanical stress.

There is an additional challenge in this fabrication that arises due to the finite aspect ratio of the vias achievable using DRIE. In a DRIE Bosch process, TSVs can be etched with aspect ratios as high as width : height = 1:10. To ensure uniform etching across the wafer, and to ensure a high yield, we use a more conservative value 1:5. Therefore, to achieve a *d* =15 *µ*m TSV, the etch depth must be *h* < 75 *µ*m. However, if we etched the delivery channels to within 75 *µ*m of the backside of the wafer, the chips became mechanically unstable and would often break during sample handling (**Fig. 2g,h**).

To overcome this challenge, in this work we incorporate a two-layer design strategy to allow small diameter vias (*d* = 15 *µ*m) to be etched without sacrificing the mechanical stability of the VLSDI chip (**Fig. 3a**). This trench approach results in mechanically stable membranes that act as effective etch stops and maintain the seal between the He cooling and the DRIE’s vacuum, leading to uniform and reproducible TSV etching over the entire 4” wafer (**Fig. 3b-d)**. To achieve this goal, we etched the via in two steps. First, we etched a trench in the delivery channel with a height *h* = 75 *µ*m and a width w = 110 *µ*m. Subsequently, the vias are etched within the trench with a height *h* = 85*µ*m and diameter *d* = 15 *µ*m to connect to the backside layer.

**Figure 3.**
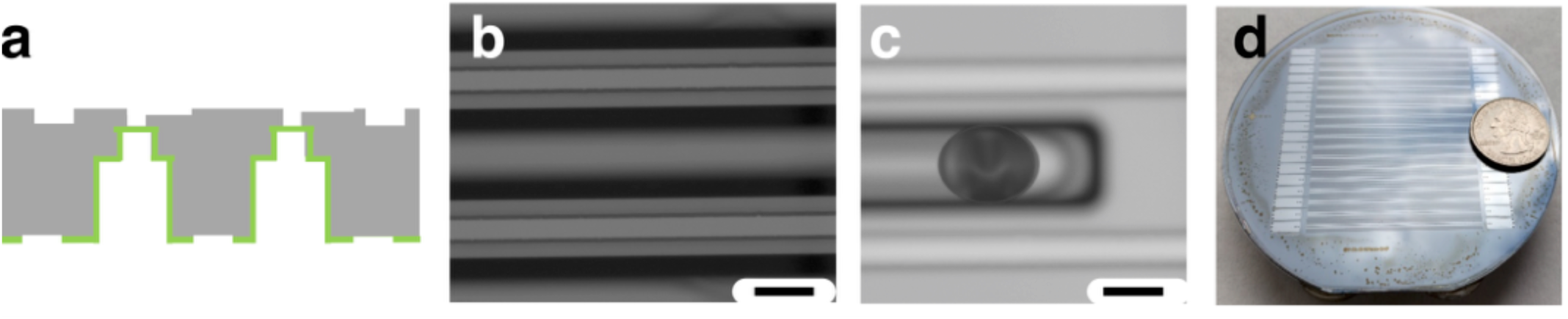
**a.** Schematic image showing our final robust fabrication strategy, including both the SiO_2_ etch-stop, stress relief, and trenches in the delivery channel. **b,c** Images show the stress relieved oxide layer and a successful *via* etch. Scale bars 200 *µ*m. (**d**) Photograph shows successfully fabricated and bonded 20k-VLSDI chip. The yield using this technique is 100% (*M* = 16 chips).

### VLSDI design for High Throughput and Low Droplet Polydispersity

The design principals and ladder geometry of our VLSDI chip have been described in detail previously. ^5,6,20^ Briefly, the main microfluidic design goals of our ultra-large-scale parallelization device is, **1.** to evenly distribute both dispersed and continuous phase fluids to each of densely packed *N* microfluidic droplet generators, **2.** to maximize the number of microfluidic droplet generators *N* that can be packed per unit area, and **3.** to maximize the generation of uniform droplets from each of these *N* droplet generators at the highest possible flow rate. These design goals and the physics of multiphase flows provide trade off relationships that guide our ultra large scale parallelization design strategies. By satisfying these design considerations, we have designed a chip that consists of 20,160 microfluidic droplet generators arranged in a 36 × 560 rectangular array with a total foot print of 6.28 × 4.93 cm^2^, with each generator having a footprint of 80 *µ*m × 1.6 mm.

The flow focusing droplet generators consist of a high aspect ratio flow resistor to ensure even distribution of flow across all 20,160 devices and a flow focusing droplet generator designed to remain in the dripping regime, by reducing the capillary number (Ca) of the continuous phase and the Weber (We) of the dispersed phase at high volumetric flow rates (**Fig. 4a-c**).^20^ The dimensions of the droplet generators are shown in **Fig. 4a-g**. The width of the flow resistors *w* = 8 *µ*m are less than the height, and thus allow the resistance to have a cubic dependence on the flow resistor’s width *R* ∝1/(hw^3^), and not the height of the entire droplet generator layer, decoupling each individual device’s flow resistance from its maximum flow rate so that droplet break-up remains in the dripping regime.^20^ The delivery channels have dimensions *w* = 0.43 mm, *h* = 0.37 mm, *l* = 54 mm (**Fig. 4h-j**). The supply channels, which supply fluid to the delivery channels, have dimensions *w* = 2.2 mm, *h* = 0.38 mm, *l* = 66 mm (**Fig 4l-o**). The trenches in the delivery channel have a *w* = 110 *µ*m and *h* = 75 *µ*m and the vias *d* = 15 *µ*m and *h* = 85 *µ*m (**Fig. 4j-k**). The underpasses, which allow the output to cross the supply lines, have the dimensions *w* = 3 mm, *l* = 7.8 mm, and *h* = 30 *µ*m (**Fig. 4l and Fig. 4o**).

**Figure 4.**
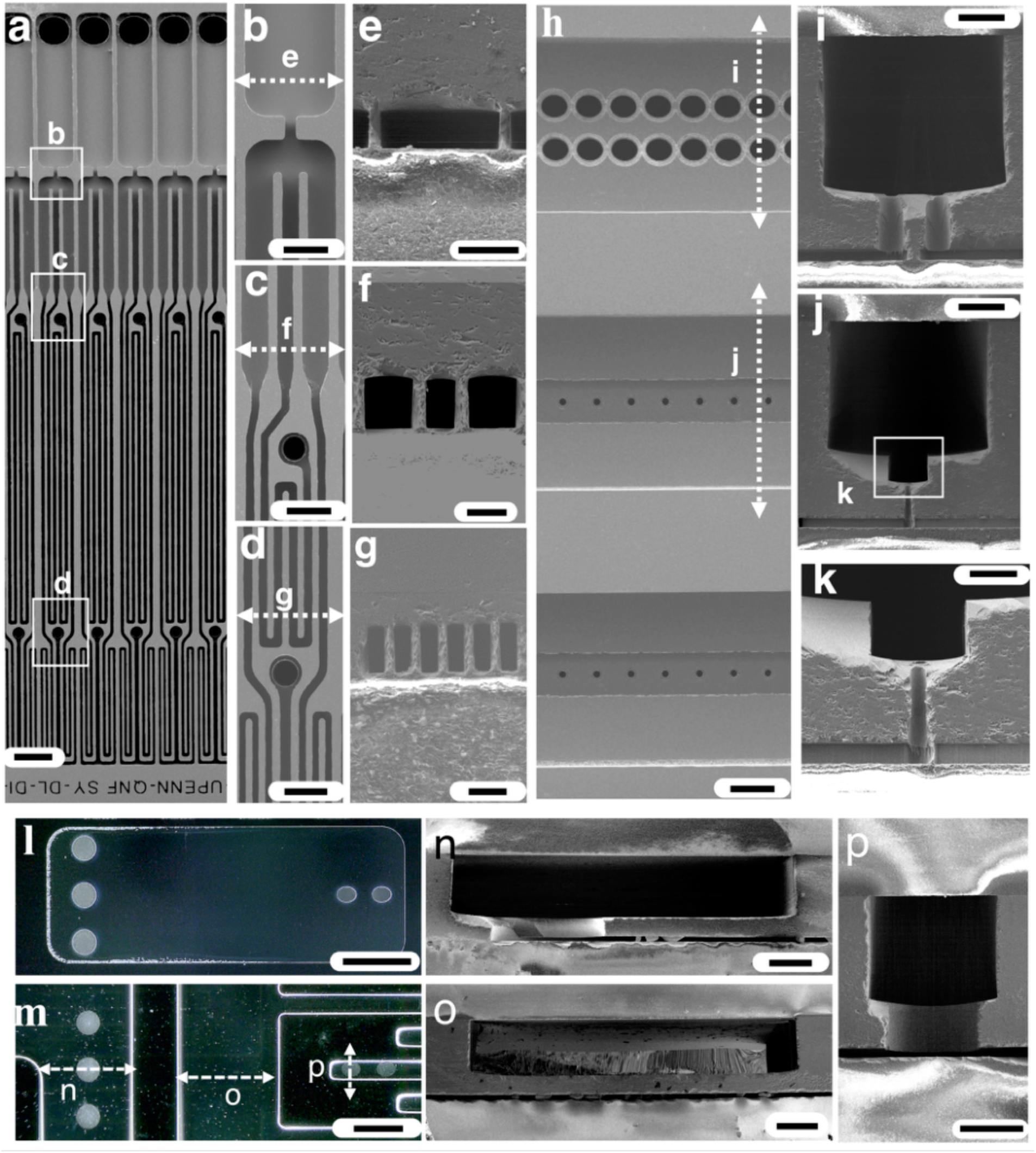
Scanning electron microscope images of the 20k-VLSDI chip. **a.** Top view of the chip.. Scale bar 80 *µ*m. Images **b**, **c** and **d** shows magnified views of **a**. Scale bars 40 *µ*m, 40 *µ*m, 33 *µ*m, respectively**. e**, **f**, **g** show cross-sections of FFG’s at positions marked in **b**, **c** and **d**. Scale bars 40 *µ*m, 20 *µ*m, 33 *µ*m, respectively. **h.** Back view of the FFGs. Scale bar 107 *µ*m. **i**, **j** show the cross-sections of delivery channels at locations marked in **h**. Scale bars 107 *µ*m and 107 *µ*m, respectively. **k.** Magnified view of the trench, via and ffg cross-sections. Scale bar 55 *µ*m. **l**, **m** show optical micrographs of top view and bottom view at underpass locations in 20k-VLSDI chip, respectively. Scale bars 1.45 mm and 1.1 mm, respectively. **n**, **o**, **p.** Micrographs show cross-section SEM micrographs at positions marked in image **m**. The Scale bars 250 *µ*m, 250 *µ*m, and 215 *µ*m.

### Droplet Generation

We first evaluated these devices by generating hexadecane droplets in water (2 wt% Tween 80). We confirmed that at all flow rates droplets are generated in every one of the 20,160 droplet generators (**Supplementary Movie S1**, **S2 and S3**). We found that our device transitioned from making uniform droplets to polydisperse droplets at a critical flow rate ϕ_dmax_ = 5.3 L/hr, resulting in a maximum throughput of 1 trillion droplets / hour (**Fig. 5a-f**). When the device was in the dripping regime, the droplets were highly monodispersed (CV <5%) and at flow rates where the droplet generator were in the jetting regime, the droplets became highly polydisperse (CV »5%). We further tested the mass production of oil-in-water emulsion by changing the dispersed phase flow rate over the range of ϕ_d_ =1.0 L/hr (20 PSI) to 5.3 L/hr (90 PSI) and the continuous aqueous phase over the range of ϕ_c_ = 1.9 L/hr (22 PSI) to 16.2 L/hr (115 PSI). By doing so, the average droplet size could be controlled over a range of *d* = 21 – 28 μm (**Fig. 6a,b**). The droplets were highly monodisperse at all flow rates, with a coefficient of variation CV < 5% (**Fig. 6c-d**).

**Figure 5.**
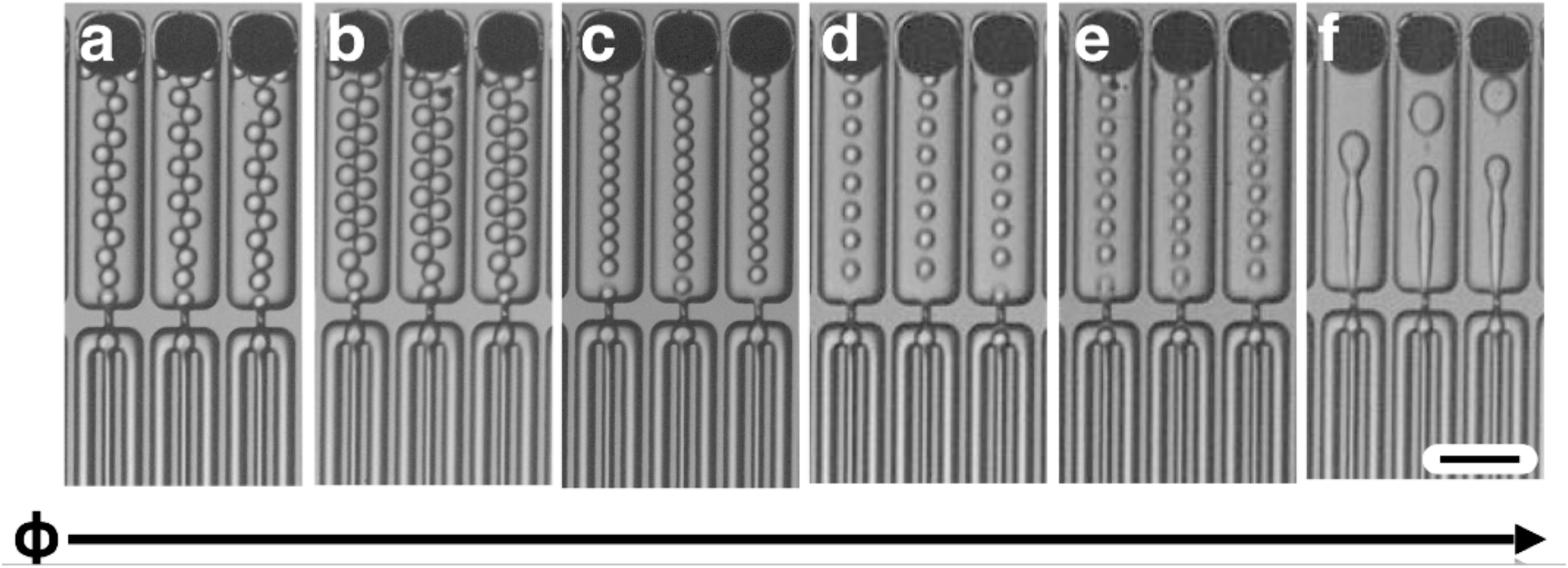
High Throughput Microdroplet Generation. **a-f.** Optical micrographs show the dripping to jetting transition in our 20k VLSDI chip for the production of hexadecane in water droplets. The maximum flow rate achieved in this device in the dripping regime was *ϕ*_c_ = 16.5 L/hr (115 PSI) and *ϕ*_d_ = 5.1 L/hr (90 PSI). The flow rates tested were (**a**) *ϕ*_c_ = 1.9 L/hr and *ϕ*_d_ = 0.9 L/hr, (**b**) *ϕ*_c_ = 1.2 L/hr and *ϕ*_d_ = 1.1 L/hr (**c**) *ϕ*_c_ = 4.5 L/hr and *ϕ*_d_ = 1.5 L/hr, (**d**) *ϕ*_c_ = 10.2 L/hr and *ϕ*_d_ = 2.3 L/hr, (**e**) *ϕ*_c_ = 16.5 L/hr and *ϕ*_d_ = 5.1 L/hr and (**f**) ϕ_c_ = 16.5 L/hr and *ϕ*_d_ = 6 L/hr Scale bar: 80 *µ*m.

**Figure 6.**
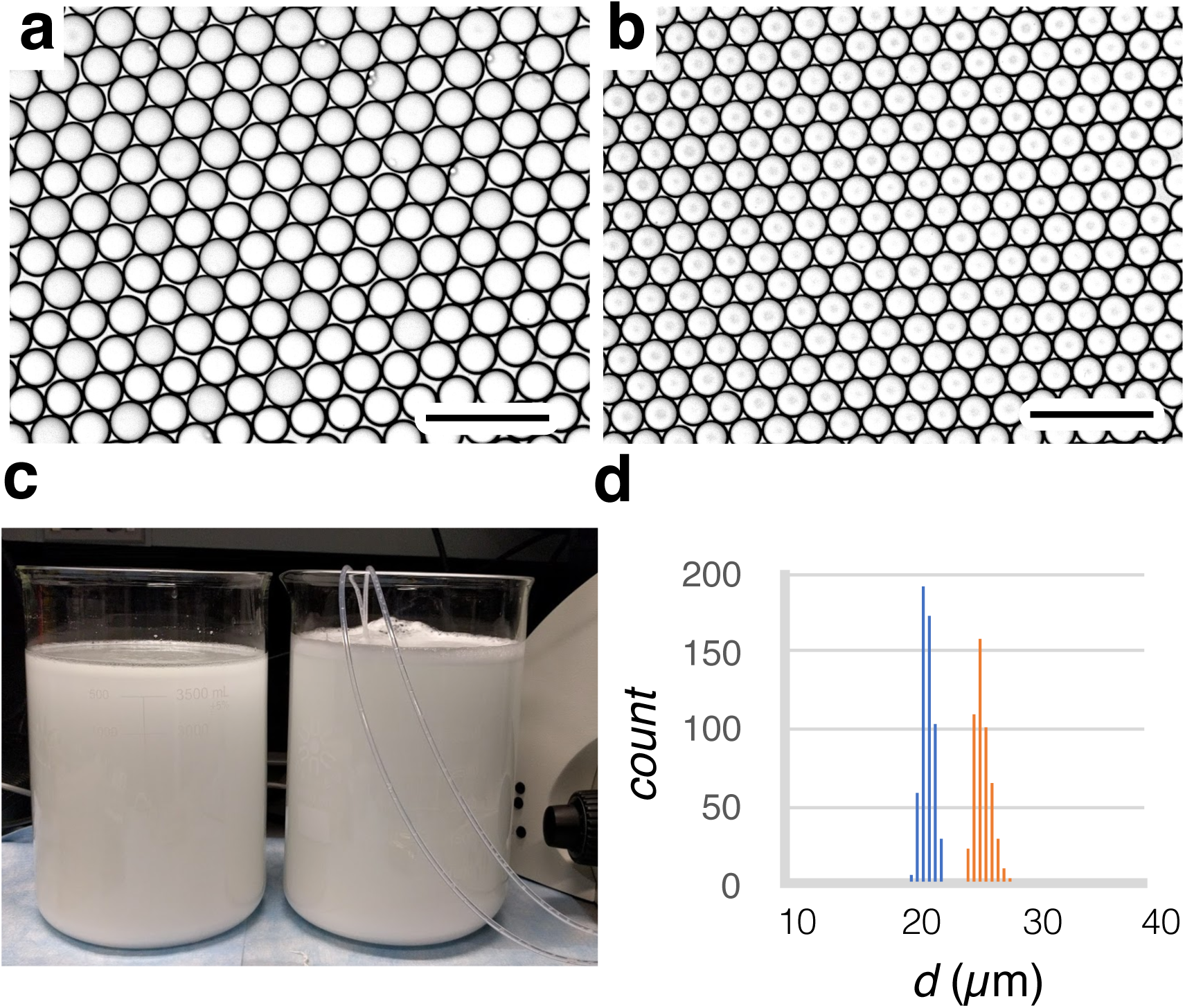
Monodisperse droplets at high throughput. **a.** Optical micrographs of *d* = 27 *µ*m hexadecane droplets in oil produced by the 20k VLSDI chip at ϕ_c_ = 1.2 L/hr and ϕ_d_ = 1.1 L/hr. **b.** Optical micrographs of *d* = 21 *µ*m hexadecane droplets in water produced by the 20k-VLSDI chip at ϕ_c_ = 16.5 L/hr and ϕ_d_ = 5.1 L/hr. **c.** 8 L of emulsion collected by running our chip for 20 minutes with *ϕ*_c_ = 16.5 L/hr and *ϕ*_d_ = 5.1 L/hr. **d.** histogram of droplets in image (**b**). For both emulsions, CV< 5%.

### Large-Scale Manufacturing of Polymer Microparticles

To demonstrate the utility of this approach in materials synthesis, we generate polycaprolactone (PCL) solid microparticles as small as *d*_p_ = 5.3 *µ*m and as large as 9.0 *µ*m, at a production rate as high as 60 grams/hr, and a CV < 5%. PCL is a biodegradable material that is approved by the United States Food and Drug Administration, and used as an injectable drug delivery system.^33^ Emulsion templates for the solid particles were generated on our chip using a dispersed phase of dichloromethane (DCM) with 4 wt% PCL. The continuous phase was deionized water with 2 wt/vol% of polyvinyl alcohol (PVA) (**Fig. 7a**, **Supplementary Movie S4**). These emulsion templates were generated on our chip at high throughput (**Fig. 7b**), collected, and processed using roto-evaporation and lyophilization. After the DCM was extracted, spherical, highly monodispersed solid PCL polymer particles were observed (**Fig. 7c,d**). To demonstrate the VLSDI’s capability to produce highly monodispersed particles with a determined particle diameter, we generated three particle formulations.(**Fig. 7e-g**) We generated particles with a diameter *d*_p_ = 9.0 *µ*m (CV = 4.0%), *d*_p_ = 7.6 *µ*m (CV = 4.3%), and *d*_p_ = 5.3 *µ*m (CV = 4.6%).(**Fig. 7h**) These particles were generated using emulsion templates produced using the following flow conditions, respectively, ϕ_c_ = 2.9 L/hr, ϕ_d_ =0.7 L/hr (35 grams/hour PCL), ϕ_c_ = 1.9 L/hr, ϕ_d_ =0.4 L/hr (21 grams/hour PCL), and ϕ_c_ = 7.7 L/ hr, ϕ_d_ =1.2 L/hr (60 grams/hour PCL), respectively.

**Figure 7.**
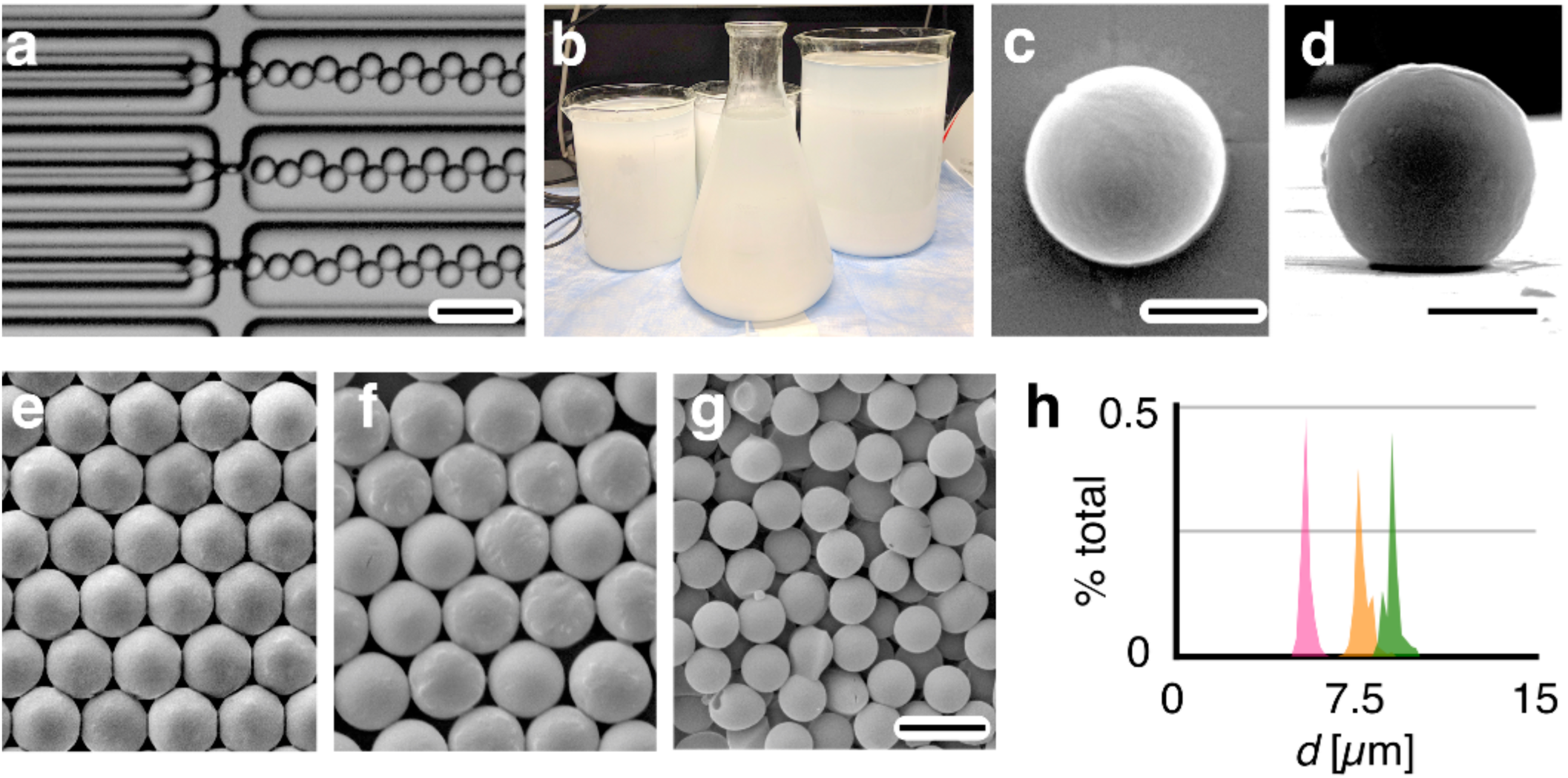
High throughput generation of solid polymer microparticles. **a.** Optical micrograph of droplet generators producing emulsion templates, with a dispersed phase of 4 wt% PCL of poly-caprolactone (PCL) in dichloromethane (DCM) and a continuous phase of 2 wt% of polyvinyl alcohol (PVA) in water. **b.** 9.0 L of emulsion collected by running our chip for an hour swith *ϕ*_c_ = 7.7 L/ hr and *ϕ*_d_ = 1.2 L/hr. Top view (**c.**) and side view (**d.**) of a solid PCL microparticle. For both images scale bar: 2.8 μm. **e-g.** SEM micrographs of three different formulations of PCL microparticles with different diameters d_p_. **h.** Histograms of the particle diameters: *d*_p_ = 9.0 *µ*m (CV = 4.0%) (green), *d*_p_ = 7.6 *µ*m (CV = 4.3%) (orange), and *d*_p_ = 5.3 *µ*m (CV = 4.6%) (pink) respectively.

## Discussion

In summary, we have developed strategies for the robust microfabrication of three-dimensional microchannels in Silicon to create highly parallelized microfluidic devices. We demonstrated the utility of these fabrication strategies by developing a VLSDI chip that incorporates 20,160 flow focusing droplet generators onto a single 4” Silicon wafer. Our microfabrication strategy allows > 50,000 TSVs to be incorporated on a single 4” wafer, with diameters as small as 15 *µ*m, with a 100% yield. Although we focus on the production of oil in water emulsions in this work, the VLSDI’s modular design enabled by the incorporation of flow resistors that decouple the design requirements for parallelization and the design of each individual microfluidic device allow for more complex designs to be parallelized for fabrication of solid polymer microparticles, multiple emulsions, micro-fibers, and nanomaterials.^12,19,33,34^

Etching of TSVs in Silicon has received much attention outside of microfluidics for its applications in three dimensional die stacking, memory stacking, CMOS, and Lab/System on a chip application.^24^ Etching of TSVs in Silicon has been done using either laser machining, wet etching, or DRIE. Laser machining is a serial process and therefore not practical with wafer-scale processes that require > 50,000 vias. Wet etching of Silicon using KOH or TMAH is limited because isotropic etches lead to larger footprint devices, because the via diameter and the via depth are coupled.^35^ DRIE, which is a parallel process, has become an industry standard to etch vias in Silicon wafers with sharp sidewalls. The strategies detailed in this manuscript can potentially be used in CMOS and MEMS fields, beyond its microfluidics applications.

## Materials and Methods

### Step-by-Step Fabrication of Microfluidic Devices

Our device has six mask layers (**Fig. SI 1-SI 6**), including layers for delivery channels (Layer 1), trenches (Layer 2), oxide mechanical stress relief (Layer 3), under-pass channels (Layer 4), through Silicon vias (Layer 5) and droplet generator channels (Layer 6). To fabricate these six layers, we produced six photomasks that are prepared on chrome-coated soda lime glass (AZ1500) using a Heidelberg 66 plus mask writer and a 10 mm write head. After exposure, all are developed in MF 319 developer for 1 minute and in chrome etchant for a minute and then the remaining photoresist is removed by immersion in 1165 developer at 60°C with sonication for 5 minutes. To fabricate the VLSDI chip, the layers are lithographically patterned, developed, and the channels are etched using deep reactive ion etching (SPTS Rapier Si DRIE). For all layers, S1805 photoresist is mixed with acetone (1:8) and the resist is spray coated to its required thickness in Suss Microtech spray coater. Spray coating is used rather than spin coating due to the deep high aspect ratio features on our chip, which would not be uniformly coated with spin coating. After each fabrication step, the wafers are cleaned using a Spin Rinse Dryer (SRD) after developing the photoresist and before DRIE.

The fabrication steps are carried out as described below, and are shown schematically in **Fig. 8a-g** and as electron micrographs in **Fig. 8h**. For the first layer (**Fig. 8 a**), we first coat the wafer with 12 *µ*m of spray-coated photoresist, soft baked on a hotplate at 90°C for 3 minutes, and exposed with the delivery channels (Layer-1)(**Fig. SI 1**) photomask. After exposure, the wafer is left at room temperature for 12 hours for rehydration. The wafer is then developed in MF 319 for 2 minutes and cleaned in SRD and then kept at 100°C on a hotplate for 10 minutes. The wafer is then cleaned again in a spin rinse dryer and etched in DRIE to achieve an etch depth of 370 *µ*m. The etched wafer is subsequently cleaned in acetone, isopropyl alcohol (IPA) and deionized water for 5 minutes each and in nanostrip for an hour and then cleaned in SRD. For the second layer (**FIg. 8b**), we first coat the wafer with 8 *µ*m of spray coated photoresist, soft baked at 90°C for 2 minutes, and exposed with the trench channels (Layer-2) (**Fig. SI 2**) photomask. After exposure, the wafer is left idle at room temperature for 1 hour for rehydration. The wafer is then developed in MF 319 for 2 minutes and cleaned in SRD and then kept at 100°C rehydration for 5 minutes. The wafer is cleaned again in SRD and etched in DRIE to a height of *h* = 75 *µ*m. The wafer is cleaned and kept in nanostrip for an hour. The etched wafer is then cleaned in SRD. For the third layer (**FIg. 8c**), we first deposit 6 *µ*m of PECVD oxide layer at a rate of 0.3 *µ*m per minute. After deposition of oxide layer, the wafer is cleaned in nanostrip for an hour. The wafer is coated with 8 *µ*m of spray coated photoresist, soft baked at 90°C for 2 minutes, and exposed with the oxide pattern (Layer-3) (**Fig. SI 3**) photomask. After exposure, the wafer is left idle at room temperature for 1 hour for rehydration. The wafer is then developed in MF 319 for 2 minutes, cleaned in SRD, and then kept at 115°C for 8 minutes. The wafer is cleaned again in SRD and etched in 25% HF for 1 minute to pattern oxide layers. The wafer is then cleaned and kept in nanostrip for an hour and cleaned in SRD.

**Figure 8.**
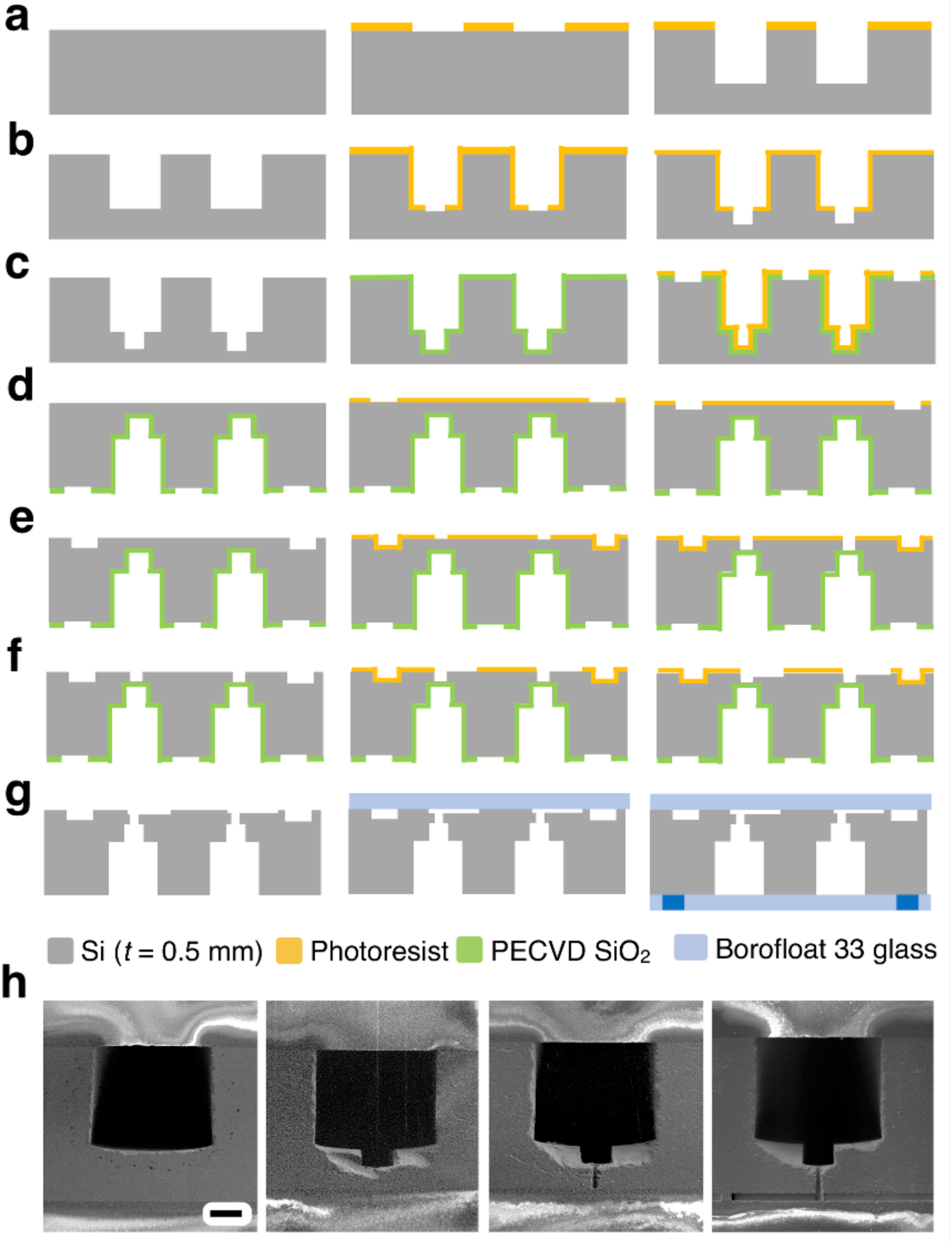
Schematic step by step description and 3D fabrication of 20k-VLSDI chip in a single 4” Silicon wafer. **a.** Fabrication of delivery channels. **b.** Fabrication of trenches inside delivery channels. **c.** Fabrication of SiO_2_ etch-stop layer. **d.** Fabrication of underpass channels in backside of wafer. **e.** Fabrication of Through Silicon Vias (TSVs). **f.** Fabrication of flow focusing generators. **g.** The 3D etched wafer and 4” Borofloat 33 glass wafers are anodically bonded to encapsulate the microfluidic channels. **h.** Scanning electron micrograph image shows the sequential etch process of fabrication in 20k-VLSDI chip. Scale bar 115 *µ*m, for ease of comparison the images are not flipped.

For the fourth layer (**Fig. 8d**), the wafer is flipped. We coat the wafer with 4 *µ*m of spray coated photoresist which is soft baked at 90°C for 2 minutes and subsequently exposed to UV with the underpass channels (Layer 4) (**Fig. SI 4**) photomask. After exposure, the wafer is left idle at room temperature for 20 minutes for rehydration. The wafer is then developed in MF 319 for 2 minutes, cleaned in SRD and kept at 100°C for 3 minutes. The wafer is subsequently cleaned again in spin rinse dryer and etched in DRIE for an etch depth of 30 *µ*m. The wafer is cleaned in acetone, IPA, water and kept in nanostrip for an hour and cleaned again in SRD. For the fifth layer (**Fig. 8e**), we spray coat the wafer with 8 *µ*m of photoresist, soft bake at 90°C for 4 minutes, and exposed with the vias (Layer 5) (**Fig. SI 5**) photomask. After exposure, the wafer is left at room temperature for 1 hour for rehydration. The wafer is then developed in MF 319 for 2 minutes, cleaned in SRD and kept at 100°C for 5 minutes. The wafer is subsequently cleaned again in spin rinse dryer and etched in DRIE for through Silicon vias. The microfabricated wafer is cleaned and kept in nanostrip for an hour. For the sixth layer (**FIg. 8f**), the wafer is coated with a monolayer of hexamethyldisilane (HMDS) in Yes Plus Oven to improve the adhesion of photoresist to the etched Silicon wafer. This step is necessary for the droplet generator layer (Layer 6) (**Fig. SI 6**), because the spacing between channels is less than 8 *µ*m and the resist can delaminate in the presence of developer or during SRD. Subsequently, 4 *µ*m of photoresist is spray coated, soft baked in an oven at 130°C for 5 minutes, and exposed to UV with the droplet maker channel (Layer 6) photomask. After exposure, the wafer is left at room temperature for 10 minutes. The wafer is then developed in MF 319 for 2 minutes and cleaned in SRD and kept at 100°C for 5 minutes. The wafer is cleaned in SRD again and etched in DRIE to a 24 *µ*m depth to form the droplet generators.

Finally, the 3D etched wafer is permanently bonded to two 4” diameter Borofloat 33 glass wafers to encapsulate the microfluidic channels (**Fig. 8g**). The 3D etched wafer is cleaned in acetone, IPA, and DI water for 5 minutes each and then in nanostrip for an hour. The 3D etched wafer and a 4 inch Borofloat 33 glass wafer are kept in piranha solution for 1 hour and immersed in deionized water for 5 minutes, and cleaned in SRD. The cleaned wafers are anodic bonded by applying 100 N force and 800 Volts for an hour in an EVG 510 anodic bonding tool. Another 4 inch Borofloat 33 glass wafer with excimer laser-drilled 1 mm holes that serve as inlets and outlets is cleaned in acetone, IPA, and DI water for 5 minutes each. The laser drilled glass wafer and the VLSDI chip are kept in piranha solution for 1 hour and then immersed in deionized water. To completely remove the piranha solution from the microchannels, a long immersion time in water is recommended. The wafers are anodic bonded, applying 100 N force and 800 Volts for an hour. Both wafers are cleaned thoroughly and handled carefully in the entire process during bonding procedure to avoid possible dust or debris that may result in weak anodic bonding (**Fig. SI 10**) that may finally result in device leakage during operation (**Fig. SI 11**). To connect the VLSDI chip to the outside world, we subsequently connect the chip to a custom-built pressure-driven flow manifold. Stainless steel compressed tube fittings (1/8”‵ tube OD) from McMaster Carr (52245K609) are bonded to the glass wafer using chemically resistant epoxy from Master Bond (EP41S-5). The epoxy is allowed to cure at room temperature for 4 days. 1/8” OD PTFE tubes were connected to the fittings. Pressure driven flow is used to conduct the experiments. Nitrogen pressure tanks were connected to 1 gallon and 3 gallon stainless steel pressure vessels (Alloy products). The 1 gallon vessel is used for dispersed phase and the 3 gallon vessel is used for continuous phase. The VLSDI chip is connected to the pressure vessels using PTFE tubings. The VLSDI chip is housed in a custom-built acrylic box and mounted onto an xyz translational stage. Inline filters (McMaster Carr: 9816K72) are used to filter debris for both the continuous and dispersed phases. An inline flow meter (McMaster Carr: 5084K23) is used to measure the flow rate of water phase. The detailed experimental setup for VLSDI chip is shown in **Fig. SI 9**.

## Acknowledgements

We thank Professor Mark Allen for his comments and inputs on our manuscript. We thank Noah Clay (Director), Meredith Metzlerm, Kyle Keenan, Hiromichi Yamamoto, Eric Johnston, David Jones, Gyuseok Kim, Charles Veith, Jarret Gilinger and all of the QNF Staff at University of Pennsylvania for their help in device fabrication. We would like to acknowledge support from The National Science Foundation (1554200), Chip Diagnostics, and The Hartwell Foundation. D.L. acknowledges the support from NSF CBET 1604536.

## Contributions

S.Y. conceived and performed all designs, fabrication, experiments, and characterization in this study, as well as prepared the manuscript and figures. D.L. and D.I. conceived and oversaw all aspects of this study, and prepared the manuscript.

