## Supplementary Information for "Robust Microfabrication of Highly Parallelized Three-Dimensional Microfluidics on Silicon"

### ELECTRONIC SUPPLEMENTARY INFORMATION

**Movies**

**Movie_S1:** Movie shows the generation of oil in water (O/W) droplets in VLSDI chip. Flowrates of Water (Qc) = 1.2 L/hr , Hexadecane (Qd)= 1.1 L/hr ;

**Movie_S2:** Movie shows the generation of oil in water (O/W) droplets at different positions in 20k-VLSDI. Oil phase: Hexadecane, Water phase: Deionized water with 2 wt% Tween 80. Flowrates of water (Qc) = 4.5 L/hr, Hexadecane (Qd) = 1.5 L/hr ;

**Movie_S3:** Movie shows the generation of oil in water (O/W) droplets at the highest flow rate tested in 20k-VLSDI chip.

**Movie_S4:** Movie shows the generation of dichloromethane (DCM) with 4 wt% polycaprolactone (PCL) suspended in water with 2 wt% polyvinyl alcohol

**Figures:**

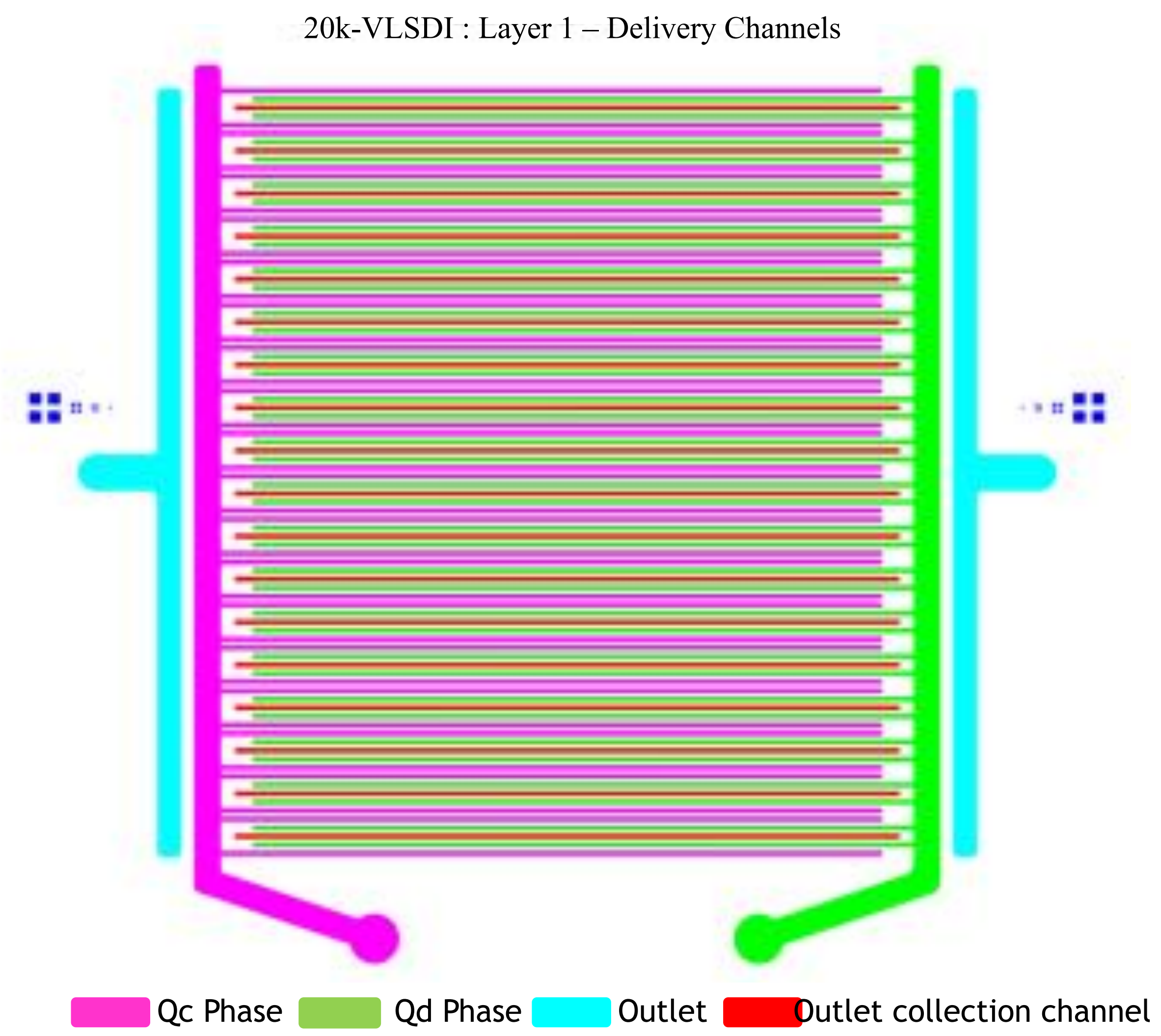


**Figure S1:** Schematic layer 1 description of 20k VLSDI chip. Qc is Continuous phase and Qd is Dispersed phase. The schematic layer shows supply channels, delivery channels, outlet collection channels and outlet channels.

20k-VLSDI : Layer 2– Trench Channels



**Figure S2:** Schematic layer 2 description of 20k VLSDI chip. Trenches  are etched in delivery channels for dispersed (Qd) and continues (Qc) phases only. Trenches are not etched in outlet collection channels.

20k-VLSDI : Layer 3 – Oxide mechanical stress relief

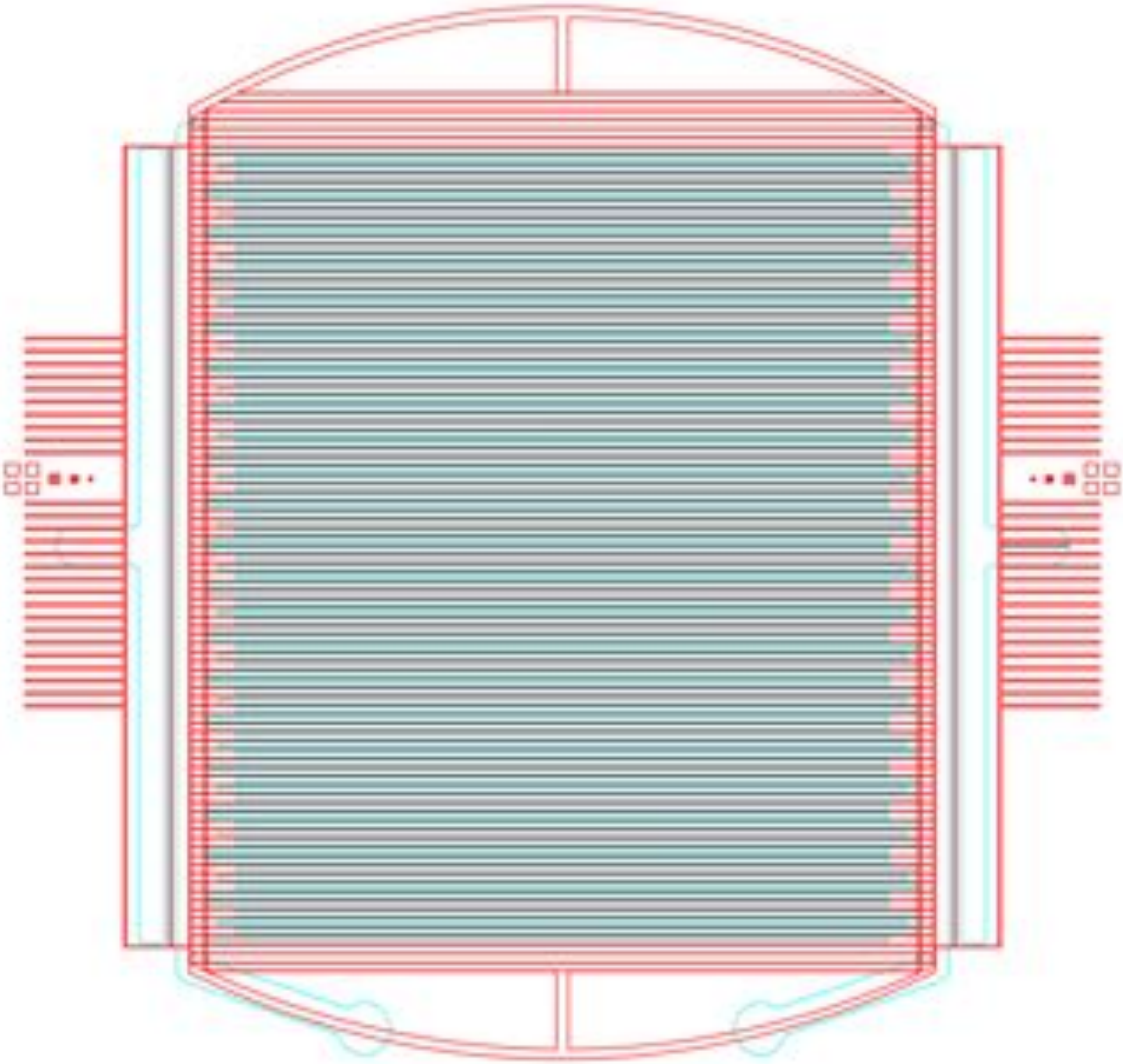

**Figure S3:** Schematic layer 3 description of 20k VLSDI chip. Layer 3 overlap on delivery channel layer (Layer-1). PECVD oxide is patterned into islands to remove the stress in as deposited oxide layer.

20k-VLSDI : Layer 4– Underpass Channels

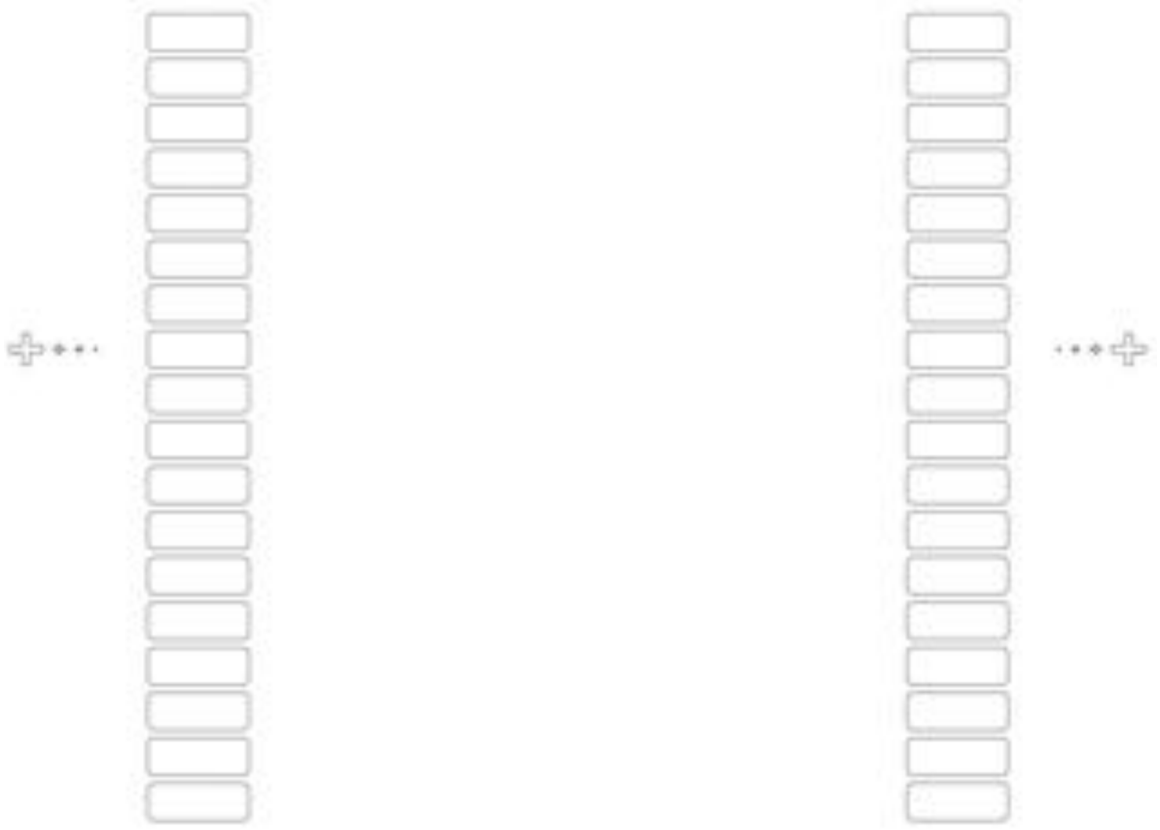

**Figure S4:** Schematic layer 4 description of 20k VLSDI chip. Underpass channels of width 2.9 mm and height 30 um and length of 7.8 mm are etched.

20k-VLSDI : Layer 5 – Through Silicon Vias

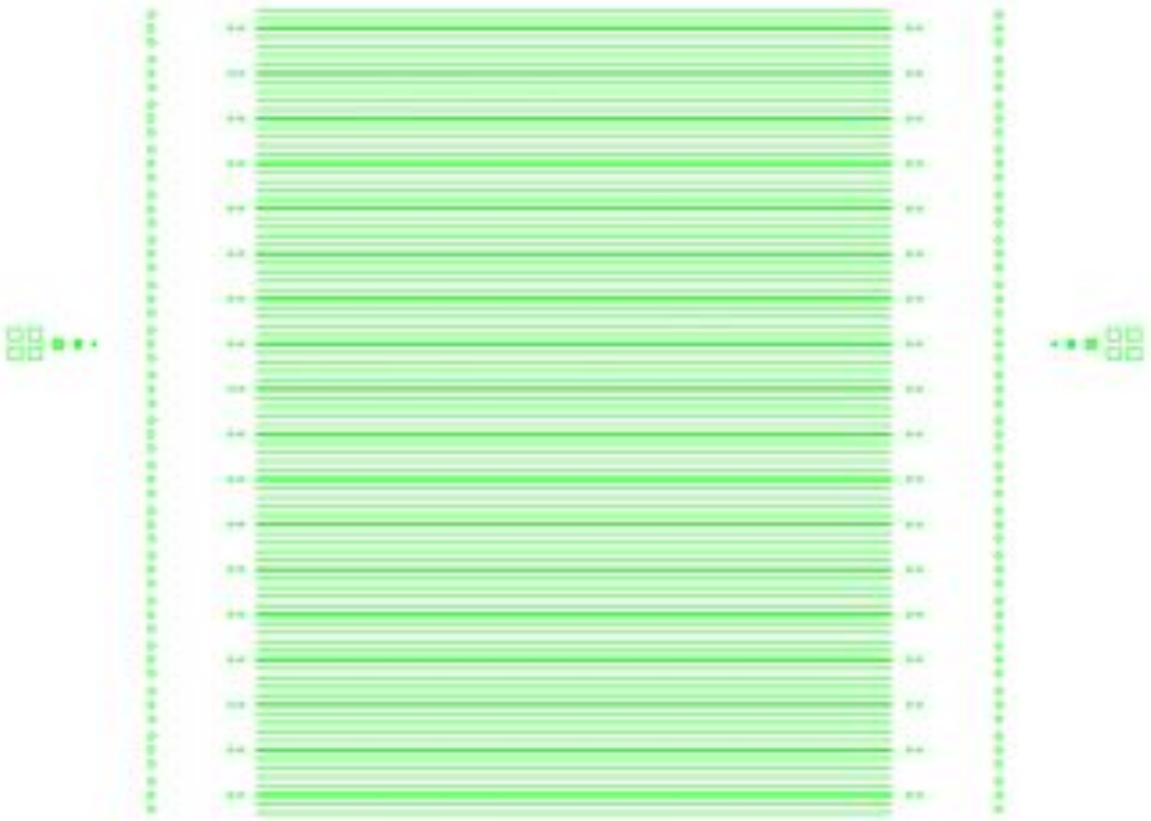

**Figure S5:** Schematic layer 5 description of 20k VLSDI chip. Approximately 60,500 Though Silicon Vias (TSV) are etched for 20,160 flow focusing droplet generators. Vias for dispersed and continuous phase are 15 um in diameters and Vias for the outlet positions for FFG's are 65 um in diameter.

20k-VLSDI : Layer 6 – Flow Focusing Droplet Generators

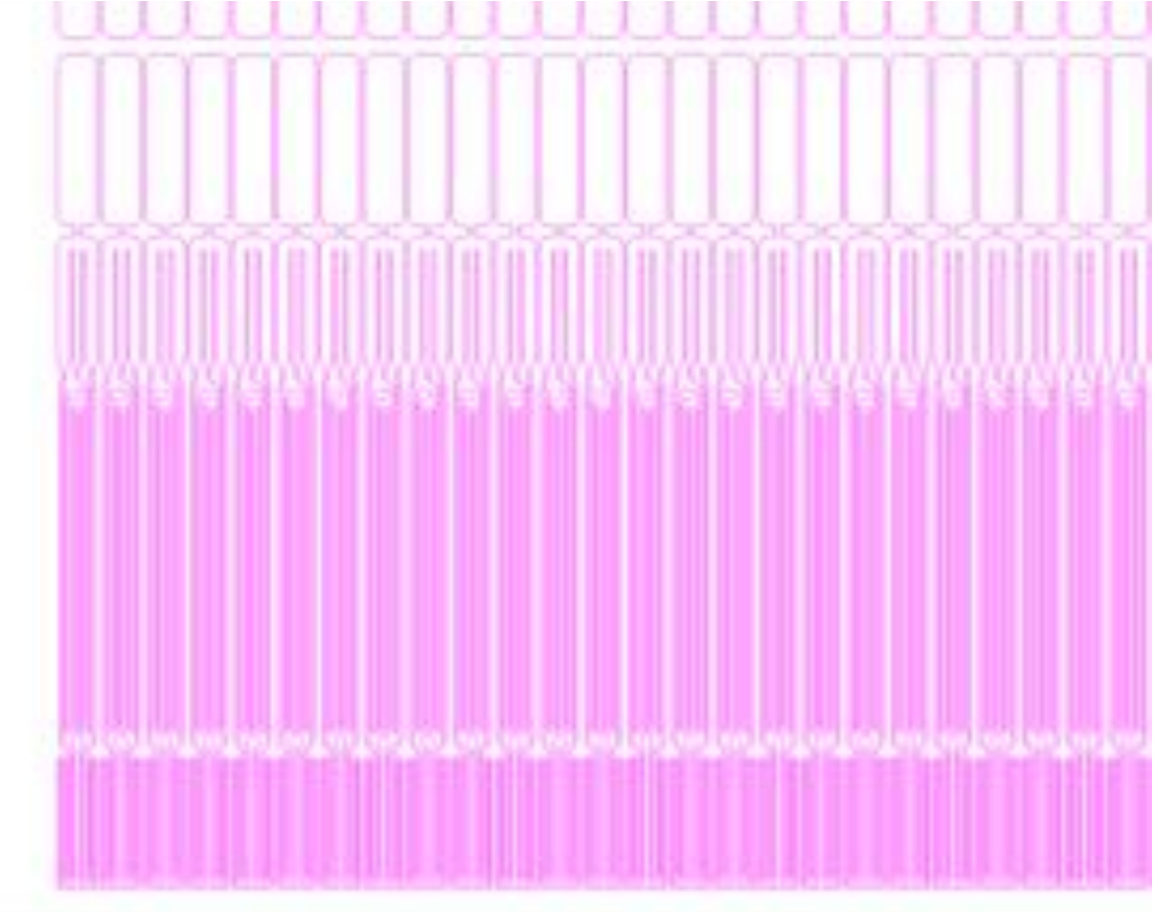

**Figure S6:** Schematic layer 6 description of 20k VLSDI chip. Flow focusing droplet generator are arranged in an array of 36 rows by 565 columns. The foot print of each device is 80 um x 1.4 mm

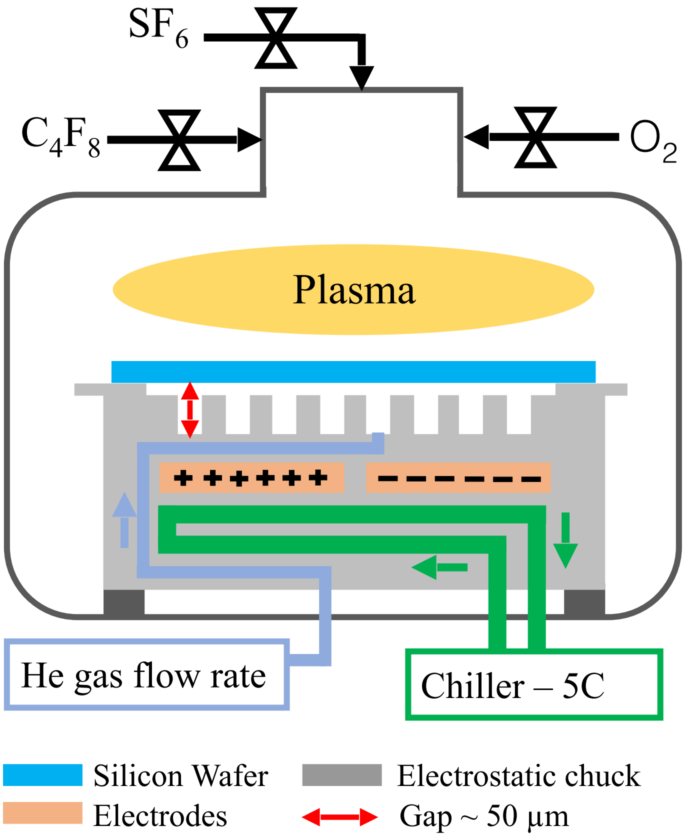


**Figure S7:** Schematic figure shows a typic DRIE chamber.

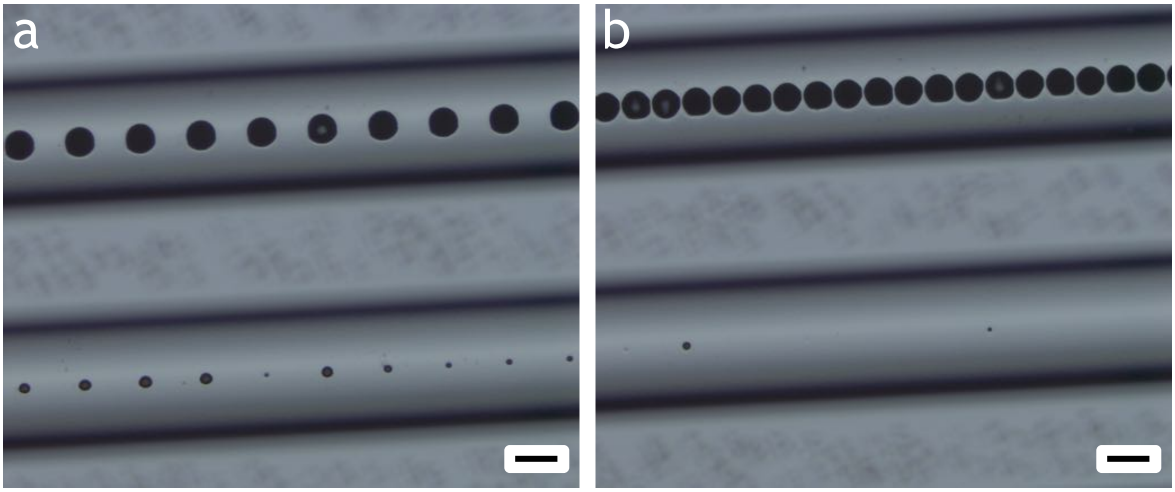


**Figure S8 : a-b.** Optical images show aspect ratio depending etching of patterns in deep reactive ion etching. Big vias etch much faster than small vias. Scale bar 200 µm.

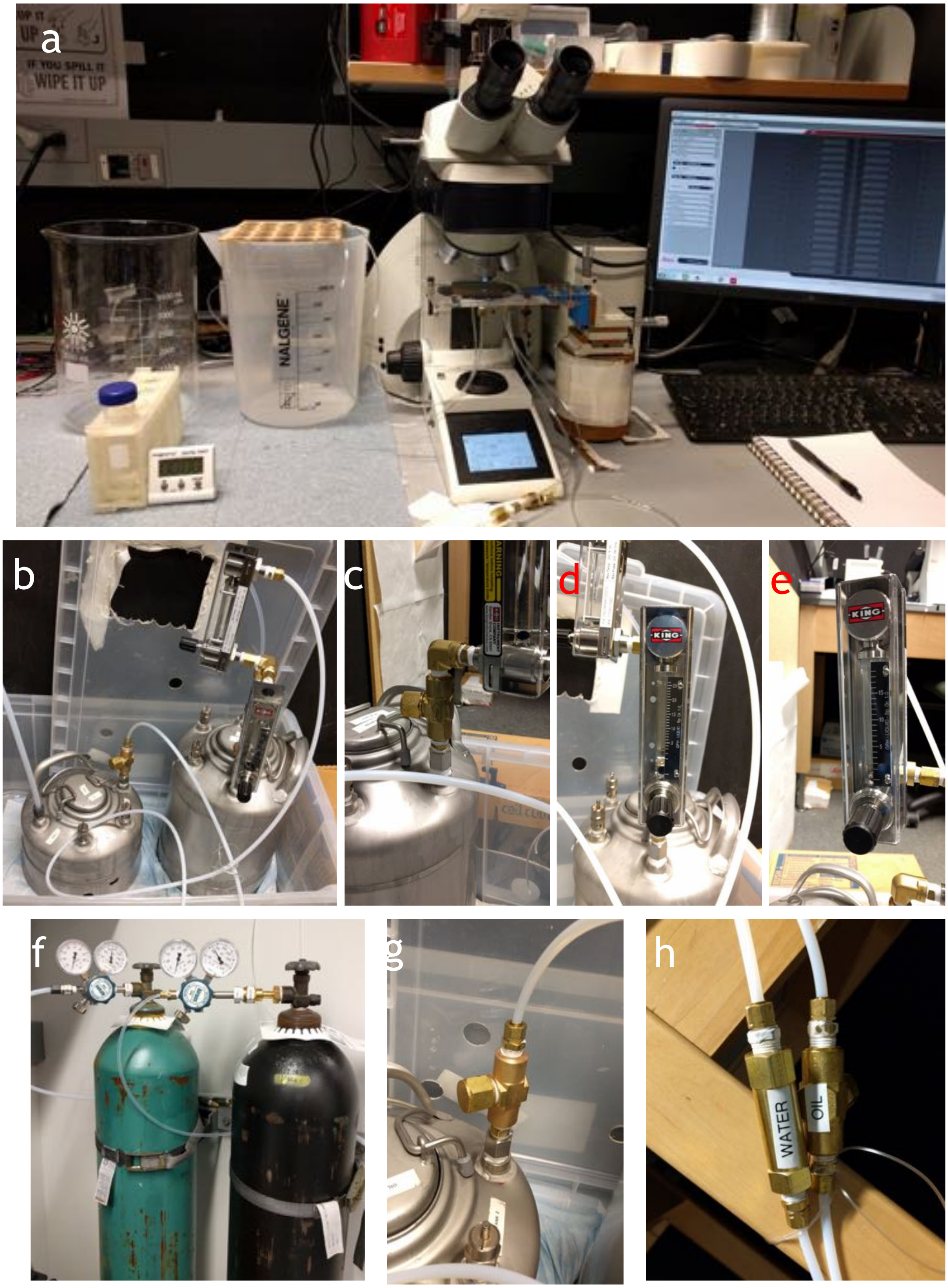


**Figure S9:** **a.** Experimental setup used to test 20k VLSDI chip. **b.** Pressure vessels used to supply fluids to chip, 1 and 3 gallon pressure vessels are used for dispersed and continuous phase fluids. **c.** Inline filter for continuous phase vessel. **d-e.** Inline flow meters for continuous phase to measure flow rates at low and high flowrates. **f.** Pressurized tanks used for pressure driven flow to vessels. **g.** Inline filter for dispersed phase vessels. **h.** Inline filter for both phases in the tubes at the positions close to the VLSDI chip.

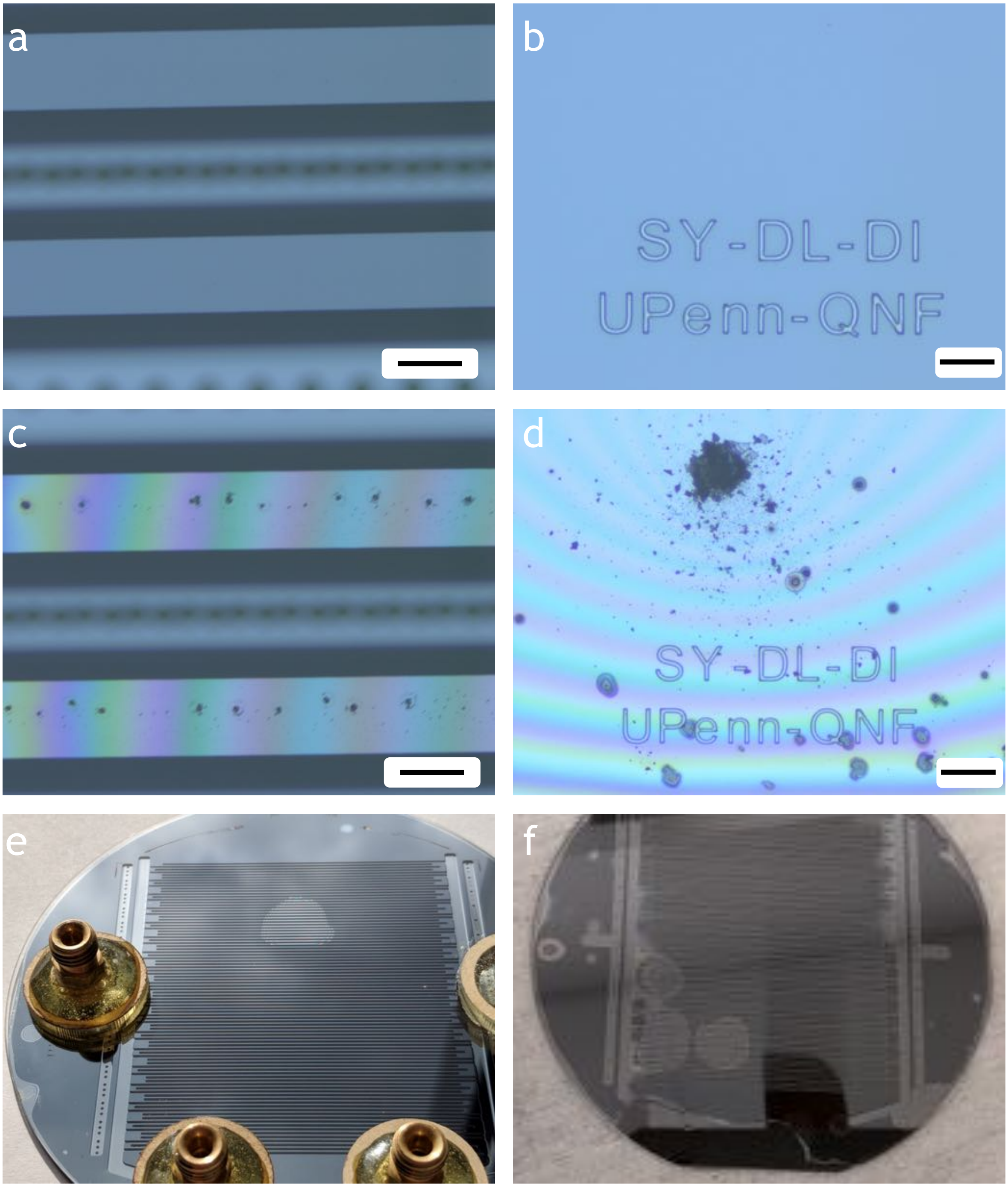


**Figure S10: a-b.** Debri free anodic bonding between glass and silicon wafer. The optical images show delivery channel side and droplet maker side of the wafer. Scale bars are 200 and 50 µm. **c-d.** Weak bonding between glass and silicon due to debris and dust particles. Scale bars are 200 and 50 µm. **e-f.** Optical images show defects in anodic bonding between glass and silicon wafers. The wafers are 4 inch in diameter. The interference pattern is a sign of defects and weak bonding between glass and silicon.

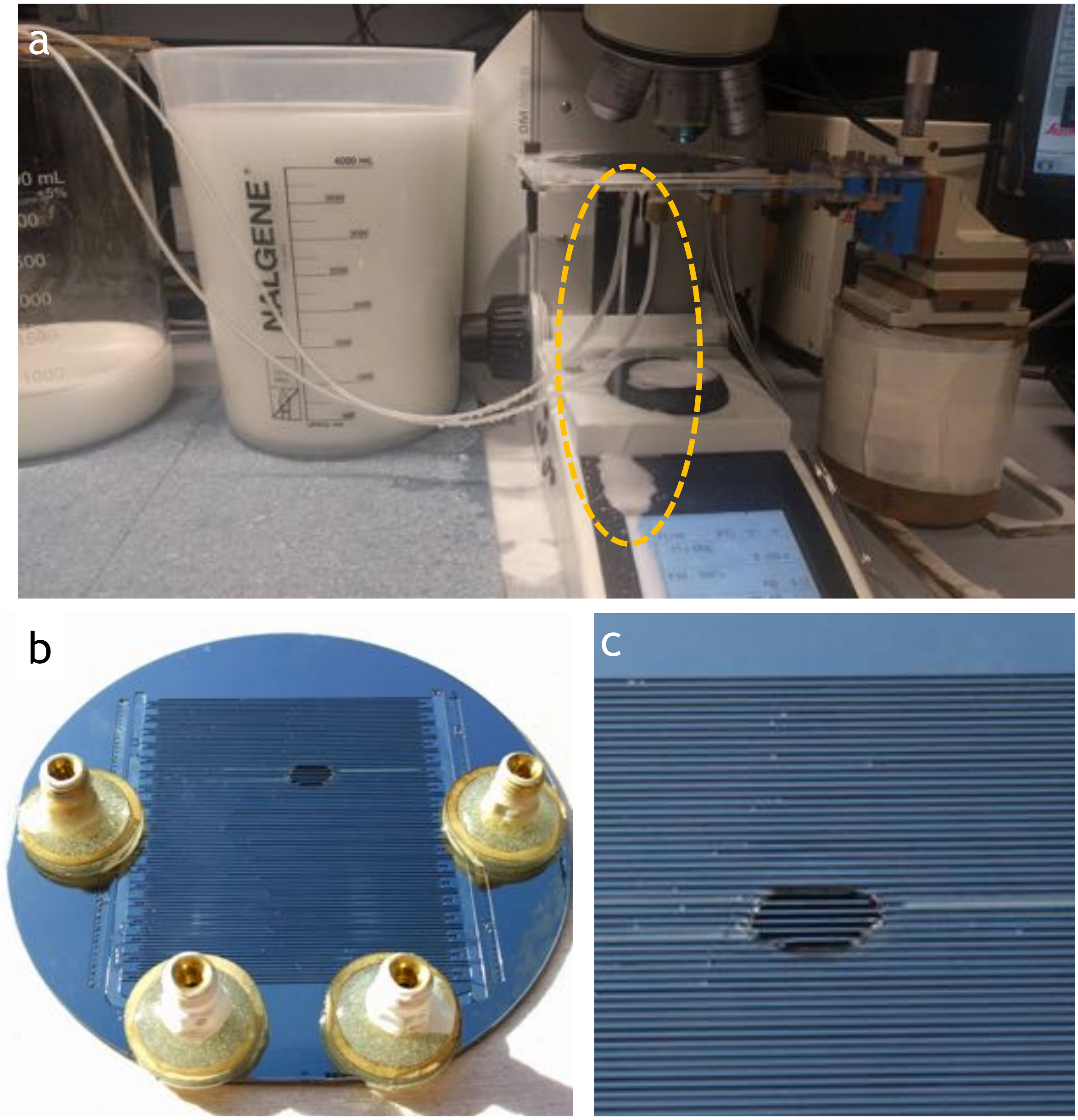


**Figure S11:** **a.** 20k-VLSDI chip leaking while operating at low flow rates due to debris and weak anodic bonding between glass and silicon. **b-c.** Optical images show broken position of the 20k-VLSDI chip while operating at low flow rates as shown in **a**.
